# A structural vista of phosducin-like PhLP2A-chaperonin TRiC cooperation during the ATP-driven folding cycle

**DOI:** 10.1101/2023.03.25.534239

**Authors:** Junsun Park, Hyunmin Kim, Daniel Gestaut, Seyeon Lim, Alexander Leitner, Judith Frydman, Soung-Hun Roh

## Abstract

Proper cellular proteostasis, essential for viability, requires a network of chaperones and cochaperones. ATP-dependent chaperonin TRiC/CCT partners with cochaperones prefoldin (PFD) and phosducin-like proteins (PhLPs) to facilitate the folding of essential eukaryotic proteins. Using cryoEM and biochemical analyses, we determine the ATP-driven cycle of TRiC-PFD-PhLP2A interaction. In the open TRiC state, PhLP2A binds to the chamber’s equator while its N-terminal H3-domain binds to the apical domains of CCT3/4, thereby displacing PFD from TRiC. ATP-induced TRiC closure rearranges the contacts of PhLP2A domains within the closed chamber. In the presence of substrate, actin and PhLP2A segregate into opposing chambers, each binding to the positively charged inner surfaces formed by CCT1/3/6/8. Notably, actin induces a conformational change in PhLP2A, causing its N-terminal helices to extend across the inter-ring interface to directly contact a hydrophobic groove in actin. Our findings reveal an ATP-driven PhLP2A structural rearrangement cycle within the TRiC chamber to facilitate folding.

**Highlights:**

- Structural analysis of TRiC-mediated folding cycle with cochaperones PhLP2A and PFD.
- The interactions of PhLP2A and PFD with TRiC are mutually exclusive.
- PhLP2A domains interact in a subunit-specific manner with the TRiC chamber.
- PhLP2A domains are rearranged in ATP-closed TRiC to contact actin across the ring interface

## Introduction

Proper protein folding and homeostasis (proteostasis), fundamental to maintaining cellular integrity, depend on the action of molecular chaperones. While *in vitro,* many molecular chaperones successfully promote the folding of non-native polypeptides, in the cell, efficient proteostasis involves the cooperation of distinct chaperones. The specificity and structural basis for such cooperation is poorly understood for most chaperone systems, and in particular for the eukaryotic chaperonin TRiC/CCT.

The ring-shaped chaperonin TCP-1 ring complex (TRiC, also called CCT) is a ~1 MDa hetero-oligomeric complex that has a double-chamber architecture, with each octameric ring formed by subunits CCT1-8 ^1^. TRiC-mediated folding depends on an ATP-driven conformational cycle, which regulates the opening and closing of a built-in lid over the central chamber of each ring ^2,3^. In the apo-state, the lid is open and substrates bind to the apical domains of specific CCT subunits ^4,5^. ATP hydrolysis leads to lid formation and substrate encapsulation in the closed central chamber, where folding occurs ^4,6–8^. Previous biophysical studies suggested stepwise folding within the chamber ^4,9^ and recent cryoEM analysis revealed that TRiC mediates a domain-wise assembly of the substrate through a directed folding pathway ^10^. TRiC plays a unique role in cellular proteostasis, as it is obligately required to facilitate the folding of ~10% of the eukaryotic proteome, including many essential proteins with a complex topology that cannot fold spontaneously or with the aid of any other chaperone ^4,11–13^.

In the cell, TRiC functions in cooperation with many cochaperones, including Prefoldin (PFD, also called GIMc) and phosducin-like proteins (PhLPs) ^14^. Prefoldin is a jellyfish-shaped hetero-oligomeric complex that binds the open state of TRiC through specific contacts with the apical domains but is released upon ATP-driven lid closure ^15^. PFD maintains substrates in a dynamic unstructured conformation ^10^ and enhances the processivity of TRiC-mediated folding by transferring substrates to the open TRiC cavity ^15^. The PhLPs are a family of ~30 kDa cytosolic proteins that have also been shown to regulate TRiC-mediated protein folding ^16^. There are five distinct PhLP homologs in humans and two homologs in yeast ^17–19^. All PhLP homologs share a similar domain organization, with a central thioredoxin-like domain ^20^ flanked by variable length flexible N- and C-terminal domains ^17^. While the precise function and mechanism of PhLPs in TRiC-mediated folding are not very clearly understood, each PhLP isoform is reported to have distinct activity and specificity for different TRiC substrates. For instance, PhLP1 assists TRiC-mediated folding and heterodimer assembly of Gβγ and Gβ5-Regulators of G protein signaling (RGS) ^21,22^. *In vitro* experiments have shown that human PhLP2A, PhLP2B, and PhLP3 can impact TRiC-mediated folding of actin and tubulin ^23,24^ and yeast PLP2, a homolog of PhLP2A stimulates the actin folding ^23^. While yeast PLP1 is dispensable for viability, yeast PLP2 is essential and interacts genetically with both tubulin and PFD-subunit Pac10 ^18,25^ Of note, PhLP2A has been mapped as a genetic determinant of a myopathy related disease ^26^.

Previous structural analyses of TRiC reported a low-resolution EM structure of PhLP bound to open TRIC ^27^ and more recently cryoEM structures of an actin-PhLP2A-TRiC complex that presumably represents a late stage intermediate after ATP hydrolysis ^28^. However, how PhLPs engage TRiC throughout the ATP-driven conformational cycle remains poorly defined. To understand the structural and mechanistic interplay between PhLP2A and TRiC throughout the ATP-driven chaperonin cycle, we used purified components to reconstitute actin folding by TRiC with its cochaperones PFD and the ubiquitous PhLP isotype PhLP2A. Here, we report a structural and biochemical analysis of the complex of TRiC and PhLP2A in distinct ATP hydrolysis states. Our study provides structural and functional insights into how PhLP2A assists TRiC-mediated substrate folding.

## RESULTS

### The architecture of PhLP2A complex with open apo-state TRiC

The complex between open, ATP-unliganded human TRiC and PhLP2A was generated using purified components and subjected to cryoEM to investigate its structure. We obtained a 3.08 Å resolution consensus map (Figure 1A, S1A-D) containing PhLP2A-dependent density within the TRiC chamber, positioned close to the equatorial region. Subsequent 3D classification and refinement revealed high-resolution features for this extra density (Figure S1B). We then fitted the crystal structure from the thioredoxin-fold domain of human PhLP2B to the density (PDB code: 3EVI) ^20^ and the model aligned well with many bulky side chains corresponding to the cryoEM density. This provides strong evidence that the extra density represents PhLP2A (Figure 1B).

**Figure 1.**
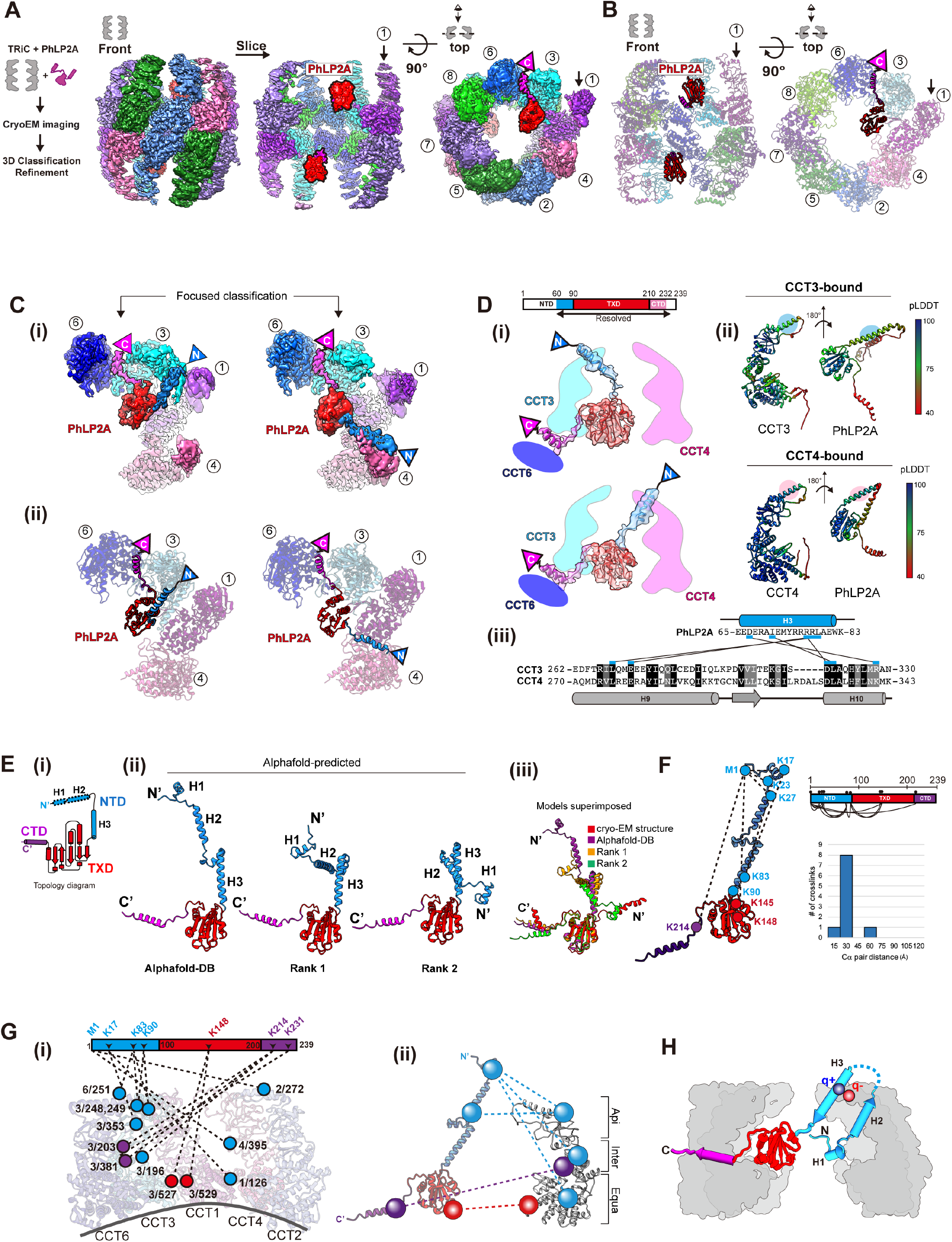
CryoEM structure of the PhLP2A-open TRiC complex. (A) (Left) Schematic of the designed experiment. (Right) Front, slice, and top view of consensus ma p of PhLP2A encapsulated in the TRiC folding chamber. Each CCT subunit and PhLP2A are color c oded as defined in the top view. (B) Atomic model of the PhLP2A-TRiC complex from the front and top view. CCT subunits and PhLP2A are color coded as in (A). (C) Two binding modes of the PhLP 2A NTD are revealed by 3D-focused classification. (i) Densities of CCT3 or CCT4 bound to PhLP2A. (ii) Atomic models of the complexes of the PhLP2A NTD and CCT3 or CCT4 using Alphafold predi ction. Each domain of PhLP2A is color-coded: NTD: dodger blue, TXD: red, CTD: dark magenta. (D) (i) Schematic diagram of PhLP2A and the CCT3/4 complex. The colored box indicates the modeled residues of PhLP2A with each domain color-coded. (ii) Alphafold prediction of each complex with a per-residue confidence score (pLDDT) diagram. (iii) Sequence alignment of H9 and H10 of CCT3 an d CCT4 at the major interaction site of PhLP2A. Predicted interactions between CCTs and PhLP2A are labeled with black lines. (E) (i) 2D Topology diagram of PhLP2A. Each domain is color coded a s indicated in (C). (ii) Three representative structures of Alphafold-predicted full-length models of PhL P2A, color-coded in a domain-wise manner and (iii) superposition of the three models with the expe rimental model. (F) Intramolecular crosslinking of PhLP2A. (Left) Intramolecular crosslinks labeled on the PhLP2A model. (Top, right) 2D schematic diagram of intramolecular crosslinks on PhLP2A. N-ter minal helices which were unresolved in cryoEM structures crosslink to all the other domains. (Botto m right) Graph of Cα pair distance. (G) Intermolecular crosslinking between PhLP2A and CCT subun its. (i) Detailed representation of the crosslinks on the model. PhLP2A crosslinks to the residues on CCT subunits of the half hemisphere of CCT2/4/1/3/6. (ii) Schematic diagram showing intermolecular crosslinks between each domain of the PhLP2A and CCT subunit. (H) Schematic representation of PhLP2A topology in an open TRiC chamber. While CTD and TXD are anchored near the equatorial domain of CCT3/6, N-terminal helices can adopt various topologies and reside inside the folding chamber.

PhLP2A consists of a coiled-coil N-terminal domain (amino acids (aa) 1-90, NTD) followed by a thioredoxin-body domain (aa 91-210, TXD) and a short C-terminal domain (aa 211-239, CTD) (Figure 1E-i). The TXD and CTD density allowed the tracing of the backbone based on sidechain densities. However, we could not detect any density attributable to the NTD. We, therefore, performed focused 3D classification on each CCT subunit (Figure S1B).

This analysis resolved an extended helical density that can be attributed to helix 3 (H3) of the NTD of PhLP2A adopting two different orientations associated with either the apical domain of CCT3 or CCT4 (Figure 1C, 1D). Next, we performed Alphafold prediction ^29^ for each possible PhLP2A-CCT subunit pair to predict residues contributing to the potential interaction. Notably, only the apical domains of CCT3 and CCT4 were predicted to form a dimeric complex with H3 of PhLP2A with good prediction scores (Figure S2A-B), in agreement with our cryoEM density maps. Therefore, we built atomic models of the complex between open TRiC and residues 63-232 of PhLP2A (Figure 1C, 1D-i), based on experimental densities as well as predicted CCT3/4 and PhLP2A contacts (Figure 1D-ii, iii, S1F, S2A-B).

To independently examine the architecture of the TRiC and PhLP2A complex, we next performed cross-linking mass spectrometry (XL-MS) analyses. The intra- and inter-molecular crosslinks observed supported and extended our cryoEM-derived model. In good agreement with our PhLP2A-TRiC, we detected many intermolecular crosslinks between PhLP2A H3 and CCT3-CCT4 as well as between the PhLP2A TXD and CTD domains to the equatorial domains of CCT3 and CCT6 respectively (Figure 1G-i, ii). While H1/H2 of PhLP2A was not identified in the cryoEM map, XL-MS identified multiple crosslinks between H1/H2 and the other PhLP2A domains. Mapping these crosslinks onto a predicted model of full-length PhLP2A (Figure 1E-F) suggests the NTD bends to place H1/H2 in the proximity of TXD (Figure 1G). Notably, we detected multiple crosslinks between the unresolved PhLP2A helices H1 and H2 with the cavity facing residues of CCT1, and CCT4. These XL-MS results together with the cryoEM structure are consistent with the PhLP2A NTD localizing inside the open TRiC chamber (Figure 1H).

### Molecular contacts between TRiC and PhLP2A in the open conformation

The cryoEM-derived model revealed that domain-specific molecular contacts mediate the PhLP2A-open TRiC interaction (Figure 2A-C and Movie S1). Overall, PhLP2A binds within the open TRiC chamber in an extended conformation where each of its domains engages distinct subunit-specific sites within the chaperonin. The central PhLP2A TXD domain is encapsulated in the open TRIC chamber where it is constrained through polar and hydrophobic interactions with the equatorial domains of CCT3 and CCT1 (Figure 2A-iii, Figure 2B-C, and Movie S1). The CTD engages in an interface formed by CCT3 and CCT6 through a single amphipathic helical structure with a highly hydrophobic side (Figure 2A-iv, v). This hydrophobic side of the CTD helix forms a hydrophobic zipper motif with an equatorial domain helix of CCT6 at the interface with CCT3 (Figure 2A-iv, v and Movie S1). The NTD of PhPL2A can adopt two different orientations, whereby H3 of the NTD contacts the apical regions of either CCT3 or CCT4. While the NTD is mostly negatively charged, H3 has a unique positively charged patch (residue 75-78), which displays a high degree of complementary to the negatively charged apical domains of CCT3 and CCT4 (Figure 2A-i, ii, 2B). While we could not resolve H1 and H2 of the NTD, we observe XL-MS contacts between these elements and the TRiC chamber including the equator of CCT1, intermediate domains of CCT3/4 and apical domains of CCT2/6. Of note, the intermediate domains of CCT3 and CCT4 have wide surfaces of positively charged residues, which may interact with the negatively charged H1 and H2 of the PhLP2A NTD (Figure 1G-H, 2B).

**Figure 2.**
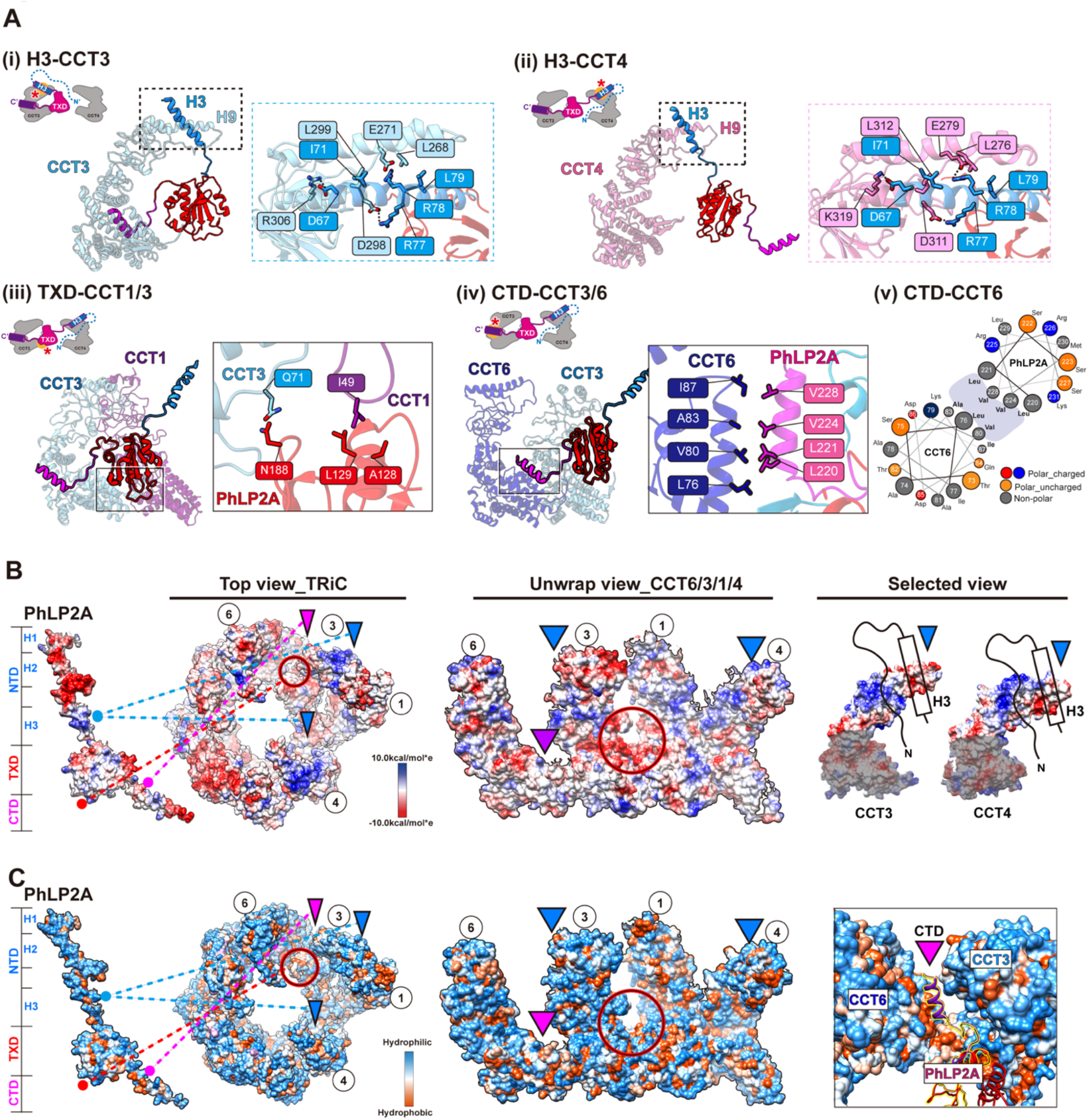
Molecular contacts between PhLP2A and open TRiC chamber. (A) Detailed interaction between PhLP2A and TRiC. (i-ii) Zoomed-in view of the binding between PhLP2A H3 and the apical domain of CCT3 or CCT4. H9 and H10 of each CCT subunit are major interaction sites; their binding modes are similar either on the topological or sequence level. (iii) Inte raction between the lower part of TXD and the nucleotide-sensing loop of CCT1. (iv) Helix-to-helix interaction between PhLP2A CTD and CCT6. (v) Helix wheel diagram on PhLP2A CTD-CCT6 showing the amphipathic nature of the helix. (B) Electrostatic surface charge of PhLP2A and open TRiC chamber. PhLP2A, the top view of TRiC, and unwrap view of the CCT4/1/3/6 half-hemisphere are dis played. Binding sites between PhLP2A and TRiC are indicated with color-coded lines and triangles (dodger blue for the NTD and dark magenta for CTD of PhLP2A). The red circle indicates the TXD binding site to TRiC. Selected views focus on the part of the apical domain where H3 of PhLP2A binds via electrostatic charge interaction. (C) Surface hydrophobicity of PhLP2A and the open TRiC chamber. Interactions between PhLP2A and TRiC are indicated in (B). While other binding sites are mainly hydrophilic, the TXD binding interface between CCT1/3 and the CTD binding crevasse between CCT3 and 6 are hydrophobic.

### PhLP2A modulates the PFD-TRiC interaction

Next, we examined the interplay between PhLP2A and cochaperone PFD, which also interacts with the open state of TRiC through the apical domains of CCT3 and CCT4 (Figure S3A). Comparing the intermolecular interfaces of CCT3/4 with PhLP2A or PFD (PDB:7WU7) showed both TRiC interactors engage in similar salt bridges with the same CCT subunits (Figure 3A). When the two models are superimposed, the contact sites of PFD and PhLP2A with CCT3 and CCT4 significantly overlap each other (Figure 3A, Figure S3A-ii). We hypothesized the steric clash between PFD and PhLP2A may lead to mutually exclusive binding, thus creating a competitive relationship between PFD and PhLP2A for TRiC binding.

**Figure 3.**
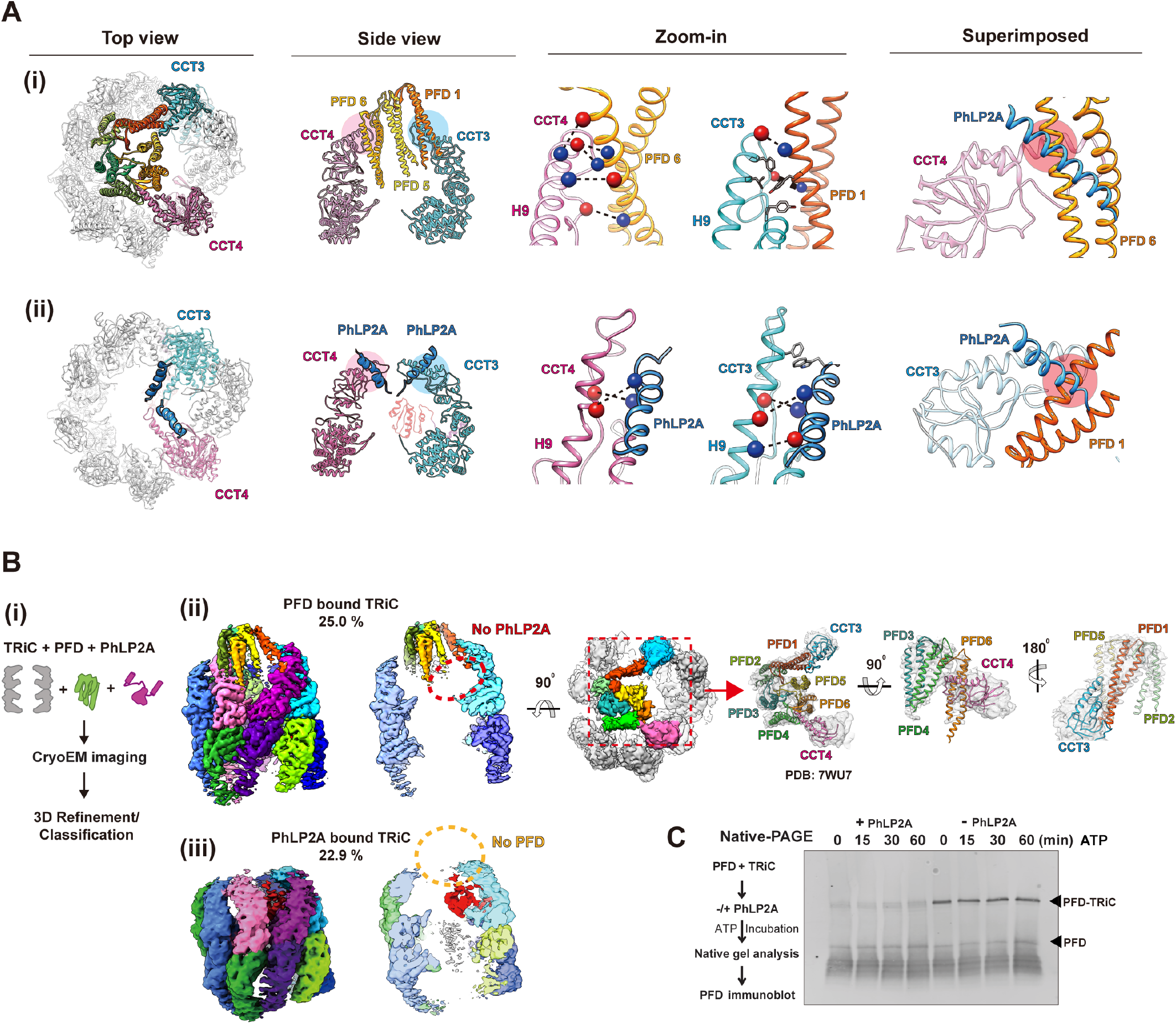
PhLP2A modulation on the PFD-TRiC network. (A) Atomic models of (i) PFD-bound TRiC (PDB:7WU7) and (ii) PhLP2A bound TRiC. CCT3, CCT4, PFD and the PhLP2A NTD are highlighted. Contacts between TRiC and PFD or PhLP2A are indicated as colored circles in the side view. Zoomed-in view of each contact between TRiC (CCT3, CCT 4) and the cochaperones (PFD, PhLP2A). Negatively charged residues are indicated by red balls while positively charged residues are indicated by blue balls. Other residues are shown as stick cartoons. PFD and PhLP2A models are superimposed on their binding sites of CCT3 or CCT4. Red circle s indicate the clash between the molecules. (B) (i) Schematic of the designed experiment. (ii, iii) 3D classification reveals PFD-bound TRiC (25.0%) and PhLP2A-bound TRiC (22.9%), respectively. The PFD-TRiC atomic model (PDB: 7WU7) fitted to the PFD-bound TRiC density. The red circle indicate s no detectable PhLP2A density while the yellow circle indicates no detectable PFD density. (C) Native gel binding assays of PFD and TRiC upon the addition of PhLP2A. Time indicates the amount o f time after ATP addition, PFD was detected by immunoblot using PFD specific antibody. PFD-TRiC complex is indicated by the arrow.

We next tested the effect of PhLP2A on PFD-TRiC binding. We generated complexes of PhLP2A and PFD with open TRiC at a TRiC:PFD:PhLP2A ratio of 1:2:~1, i.e. a two-fold ratio of PFD to TRiC and PhLP2A. When analyzed by cryoEM (Figure 3B-i), the particles displayed high heterogeneity with respect to the binding states of the cochaperones to TRiC (Figure S3B, S3C). We performed 3D-focused classification on each location of PFD and PhLP2A independently and found that TRiC particles only contained either PFD or PhLP2A (Figure 3B, S3C). This observation indicates PFD and PhLP2A binding to TRiC is mutually exclusive. To cross-validate this observation, we used Native PAGE (N-PAGE) analyses to assess if the addition of PhLP2A affects the formation of a PFD-TRiC complex (0.25 μM TRiC, 2 μM PFD-actin, 2 μM PhLP2A). When incubated with TRiC, PFD binds and forms a super-shifted complex migrating close to the TRiC band in the N-PAGE. Of note, the addition of 2 μM PhLP2A to the incubation, reduces the amount of PFD bound to TRiC, consistent with the prediction from cryoEM analysis that PhLP2A precludes PFD binding to TRiC (Figure 3C). It is worth noting that we performed additional experiments in different molar ratios in an attempt to obtain a TRiC structure with PhLP2A and PFD in each opposite chamber, but interestingly, we couldn’t get such a structure in any molar ratio.

Our analyses showing both PFD and PhLP2A associate with TRiC through an electrostatic interface with subunits CCT3/CCT4 suggest a mechanistic underpinning to orchestrate their action on a TRiC-centered network. To assess the functional relevance, we asked if these interaction interfaces are conserved in evolution. Indeed, the charged patch on CCT4 is highly conserved and its mutation impairs cellular proteostasis ^15^. Residue conservation analysis for each chaperone ^30^ indicates that residues forming the contact surface among TRiC, PFD and PhLP2A are evolutionary conserved. Particularly, PFD6 has a highly conserved positively charged surface that contacts CCT4 (Figure S3F). The charged H3 residues in PhLP2A that contact CCT4 are also highly conserved (Figure S3F). Finally, the contact areas of CCT3 and CCT4 toward PFD and PhLP2A are evolutionarily conserved (Figure S3G), as described. The evolutionary conservation of the inter-chaperone interaction sites suggests a coevolution between these three chaperones to coordinate substrate delivery and TRiC-mediated folding.

### The architecture of PhLP2A complex with ATP-induced closed state TRiC

TRiC conformation changes upon ATP hydrolysis, which induces lid closure over the central chamber ^3,6,31^. Since lid closure occludes the PFD binding site in CCT3/4 and induces the release of bound PFD from TRiC, we examined if PhLP2A forms a complex with closed TRiC. We incubated the TRiC: PhLP2A in a 1:4 molar ratio with ATP/AlFx, which generates a stably closed TRiC conformation and subjected it to cryoEM analysis (Figure 4A-i). The consensus map of closed TRiC reached 2.95 Å resolution with significant additional density inside the chamber. Subsequently, focused classification on the inner TRiC chambers identified subpopulations of PhLP2A in either one (14.0%) or both chambers (9.5%) (Figure S4A-D). The TRiC subpopulation with PhLP2A in both chambers was used to further refine the final map of the PhLP2A protein fully encapsulated inside the closed TRiC chamber at 3.24 Å resolution (Figure 4A-ii, iii). Overall, side chain details for TRiC were clearly revealed on the map allowing us to build an atomic model. For PhLP2A, we could build a refined model spanning residues 27-214 (Figure 4B-i, S4E). PhLP2A H1-2 (aa1-26) in the NTD and the CTD (aa 215-239) are not well resolved but have attributable densities that can be observed at a lower contour providing an approximate location of the NTD in proximity to CCT1 and CCT3 and of the CTD in proximity to the intermediate domains between CCT1 and CCT4 (Figure 4B-ii).

**Figure 4.**
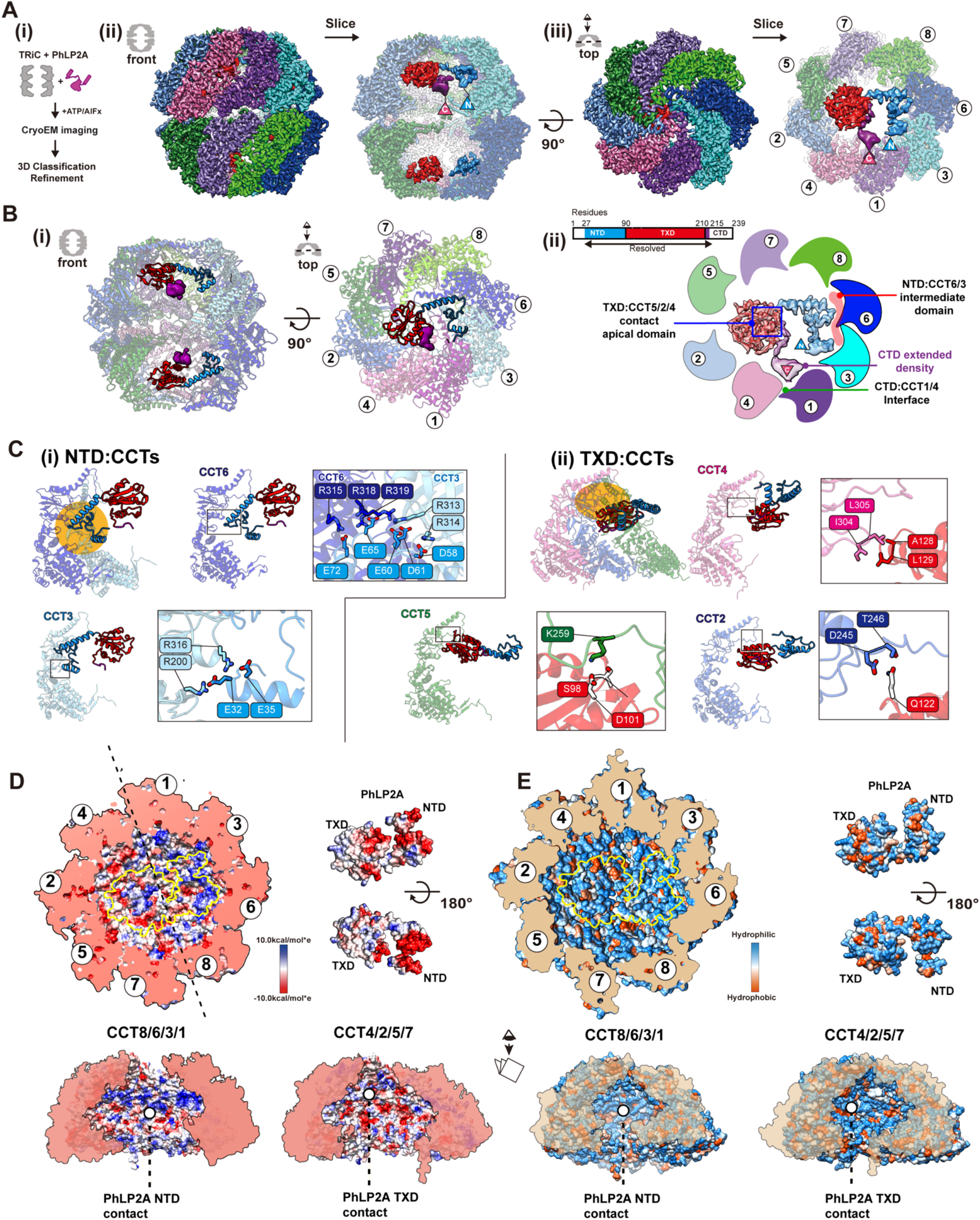
Structure of PhLP2A inside TRiC chamber after chamber closing. (A) CryoEM structure of PhLP2A inside the closed TRiC chamber after ATP hydrolysis. (i) CryoEM imaging preparation scheme. (ii) Front and slice view of the closed PhLP2A-TRiC complex. C- and N-terminus of PhLP2A are labeled in the map. (iii) Top and slice view of the closed PhLP2A-TRiC complex. Due to the varying resolution, the PhLP2A density is shown at different thresholds according to the protein domains. Note that both of the folding chambers can be occupied by PhLP2A, as in open-state TRiC. Each subunit and domain are color coded as in Figure 1. (B) (i) Model of PhLP2A inside the closed TRiC chamber. The CTD is not modeled and the lowpass filtered electron density is shown instead. (ii) Summary of binding sites between PhLP2A and the closed TRiC chamber. Compacted NTD helices interact with the intermediate domain of CCT3/6 (colored in red) and the foll owing TXD interacts with the apical domain of CCT5/2/4 (marked by a blue box). Note that although these domains are not resolved to the atomic level, the density map at the lower contour level shows the NTD close to CCT1/3 and the CTD close to the equatorial domains between CCT4/1. (C) Zoom-in view of the domain-wise interaction between PhLP2A and CCT subunits. (i) Molecular conta cts between CCT 3/6 and the NTD of PhLP2A and (ii) between CCT5/2/4 and the TXD of PhLP2A. Contact areas are indicated by a yellow circle. Residues making contacts are displayed and labeled. (D) Electrostatic surface charge showing charge complementarity between PhLP2A and the closed folding chamber. The positively charged half hemisphere of CCT1/3/6/8 provides a binding surface for the negatively charged PhLP2A NTD. (E) Hydrophobic surface charge of closed folding chamber an d PhLP2A. While the inner wall of the chamber is mostly hydrophilic and charged, part of the surface composed of CCT5/2/4 provides hydrophobic patches for PhLP2A TXD to bind on.

Remarkably, PhLP2A also interacts with the closed TRiC chamber through domain-specific contacts with specific CCT elements, albeit these are distinct from those observed in the open state. Negatively charged residues in the NTD form multiple salt bridges with positively charged residues exposed by CCT3 and CCT6 in the closed chamber, these contacts constrain the NTD to bind this region of the TRiC inner wall (Figure 4C-i and Movie S2). The TXD localizes beneath the lid region through polar and hydrophobic interactions with the lid regions of CCT5, 2, and 4 (Figure 4C-ii and Movie S2). The CTD density at the lower contour is in proximity to the interface between the intermediate domains of CCT1/4. Previous analysis recognized that the TRiC inner chamber contains an asymmetric charge distribution, with a strong positively charged inner surface contributed by subunits CCT1/3/6/8 and a strong negatively charged inner surface contributed by CCT7/5/2/4 ^1^ (Figure 4D). The strong electrostatic binding between the negatively charged NTD of PhLP2A and the positive TRiC hemisphere together with the interaction between TXD of PhLP2A and the TRiC lid segments on the other hemisphere (Figure 4D-E) establishes a diagonal binding topology of PhLP2A inside the closed TRiC chamber (Figure 4C and Movie S2).

### ATP-driven TRiC closure changes the conformation of bound PhLP2A

Our analysis shows that, unlike PFD, PhLP2A binds TRiC in both open and closed states. ATP hydrolysis leads to many changes in available TRiC interaction interfaces, including in the apical domains, the formation of an asymmetrically charged inner chamber wall, and the formation of the lid all of which produce a global reorientation of the PhLP2A domains within the TRiC chamber (Figure 5A-B and Movie S3). The transition from open to closed TRiC triggers a ~50 Å movement of the negatively charged NTD H3 of PhLP2A from binding the apical domains of CCT3 and CCT4 to binding the positive inner wall of the closed TRiC at CCT3 and CCT6 intermediate domains (Figure 5A-B and Movie S3). In addition, TXD rotates ~180 degrees and moves ~30 Å upon lid closure, from equatorial contacts to CCT3-CCT6 in the open state to contacting the opposite hemisphere at the apical CCT5/2/4 closed lid segments (Figure 5A-B). Finally, the PhLP2A CTD, which is also tightly anchored to the CCT3/6 interface in open TRiC, is released in the folding chamber upon TRiC closure, with weak contact with the CCT1/4 inner wall. In conclusion, the changes in surface residues within the TRiC chamber upon lid closure induce a dramatic conformational repositioning of PhLP2A by reassortment of its domain interactions with the chamber.

**Figure 5.**
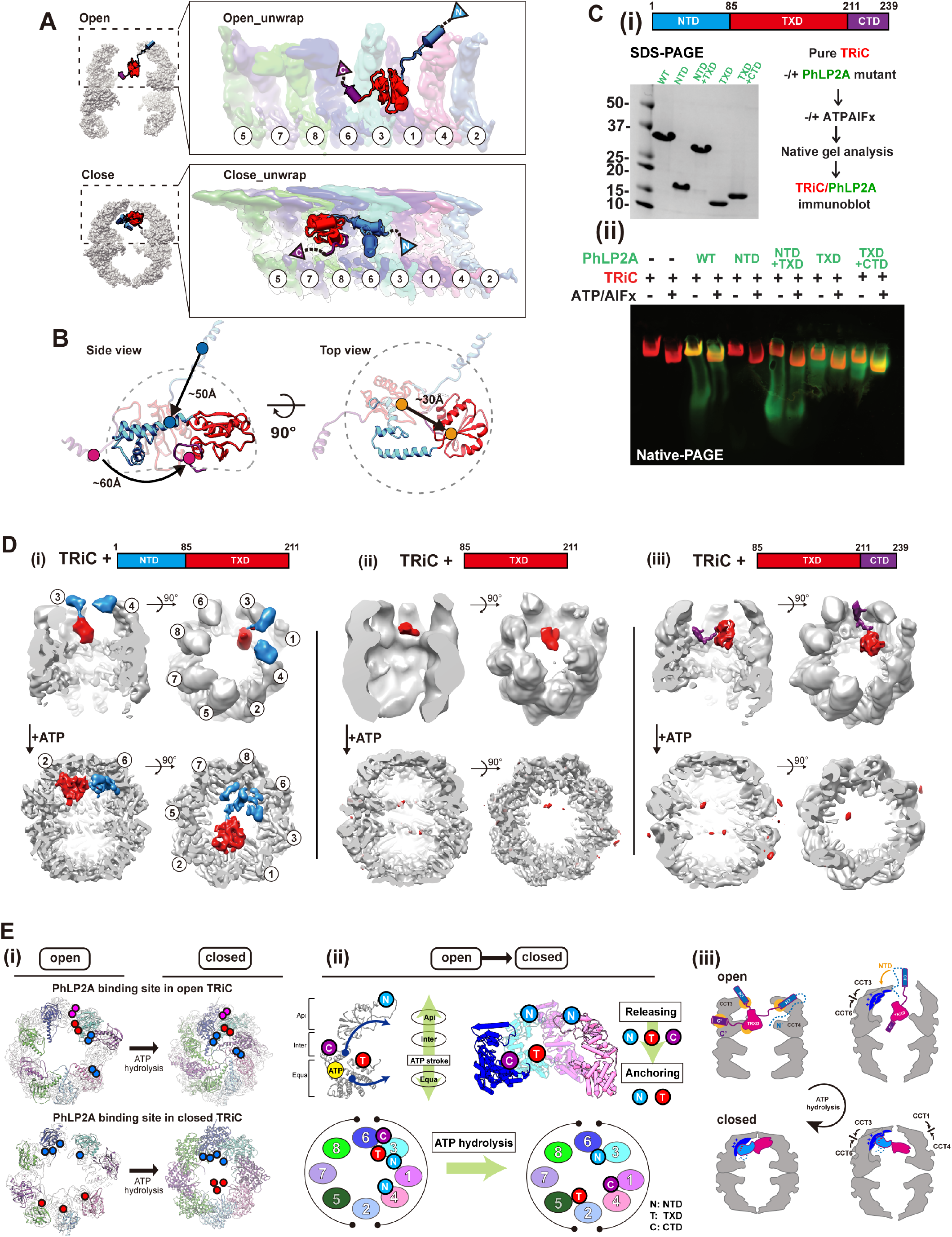
Domain-wise characteristics of PhLP2A in relationship with TRiC. (A) Global rearrangement of PhLP2A in the TRiC folding chamber in the unwrapped view. (Top) Unwrapped view of PhLP2A in which NTD is in the expanded form in TRiC in the open conformation. (Bottom) Unwrapped view of PhLP2A in the closed conformation of TRiC. Note that PhLP2A NTD is compacted and constrained. (B) Conformational and orientation changes of PhLP2A in the open or closed TRiC folding chamber. PhLP2A NTD undergoes an orientation change of about ~ 50 Å from e xtended outward to bent inward, closer to the TXD. The TXD is lifted about ~ 30 Å from the equat or to the apical contact through flipping but retains its conformation. The CTD moves ~ 60 Å followi ng the movement of TXD. (C) (i)(left) SDS-PAGE of PhLP2A wild-type (WT) and four truncated mutant constructs of PhLP2A: NTD (aa 1-84), CTD truncation (aa 1-211, NTD+TXD), TXD (aa 85-211), N TD truncation (aa 85-239, TXD+CTD). (right) Schematic of the binding assay. (ii) Binding assay between TRiC and PhLP2A truncation mutants with or without ATP/AlFx. (D) CryoEM structures of three truncated mutants inside the TRiC in the open state and after ATP hydrolysis. (i) TRiC with NTD+T XD shows attributable NTD density at the apical domain with TXD and NTD+TXD orients like WT P hLP2A in closed TRiC. (ii) TRiC with TXD: attributable density of TXD at low resolution at the equator, but no attributable density in closed TRiC. (iii) TRiC with TXD+CTD: CTD anchor shows high-res olution features like WT PhLP2A between CCT3/6 and TXD close to CCT3, but no attributable density in closed TRiC. (E) (i) Schematic diagram of PhLP2A domain-wise interaction with the TRiC cham ber in open (left) and closed conformation (right). Colored circles indicate interacting residues in ope n TRiC and closed TRiC and their movements during the ATP cycle. Each PhLP2A domain is color coded as in (C). (ii) ATP hydrolysis event in CCT subunits cascades the releasing and re-anchoring process of PhLP2A. N: NTD, T: TXD, C:CTD. (iii) Diagram of releasing of PhLP2A upon ATP-depen dent cycle of TRiC.

### Dissecting PhLP2A domain-specific interactions with open and closed TRiC

To better define the interactions of PhLP2A domains with specific TRiC subunits, we generated four domain variants: NTD (aa 1-84), TXD (aa 85-211), NTD-TXD (aa 1-211) and TXD-CTD (aa 85-239) respectively (Figure 5C-i). To assess TRiC binding, we incubated either full-length PhLP2A or four domain variants with bovine TRiC (0.5 μM TRiC, 5 μM PhLP2A variant) (Figure 5C-ii, Figure S5A). Binding was assessed by N-PAGE followed by immunoblot against the His-tag on PhLP2A to examine if the PhLP2A variants comigrated with either open or ATP-closed bovine TRiC (Figure 5C-ii, Figure S5A). While the NTD alone does not bind TRiC, the TXD alone bound TRiC in both the absence or presence of ATP/AlFx. Of note, the TXD-CTD variant increased TRiC binding over TXD alone. We conclude that the TXD is sufficient to mediate TRiC-binding but the CTD enhances this interaction. Of note, these domain variants can still be encapsulated by TRiC lid closure (Figure 5C, Figure S5A).

To further dissect the molecular determinants of TRiC binding, complexes of PhLP2A domain variants with open and closed TRiC were examined by cryoEM (Figure 5D, Figure S5B-C). For the open TRiC complexes, we identified extra density attributable to the PhLP2A variants with similar placements as identified in full-length PhLP2A. The NTD-TXD fragment in the TRiC open chamber showed similar density attributable to the NTD around the apical helices of CCT3/CCT4 but displayed higher heterogeneity around the TXD density (Figure 5D-i). TXD-only showed density close to the TRiC equator only at a lower contour level, suggesting its intrinsic affinity for the TRiC chamber depends on other PhLP2A domains (Figure 5D-ii). Indeed, TXD-CTD yielded high-resolution features like WT PhLP2A, with the CTD anchored between CCT3/CCT6 and the TXD close to CCT3 (Figure 5D-iii). The complex particle population of TXD-CTD was notably higher and the structural features were sharper compared to NTD-TXD and TXD only, indicating that CTD plays a key role in PhLP2A binding to open TRiC, consistent with N-PAGE (Figure 5C-D, Figure S5C). The addition of ATP/AlFx encapsulated all PhLP2A variants and rearranged their interactions within the closed TRiC chamber (Figure 5C). While NTD-TXD reoriented similar to full-length PhLP2A in the closed TRiC chamber, we could not observe any density attributable to either TXD or TXD-CTD in closed TRiC (Figure 5D), even though N-PAGE indicated all PhLP2A variants remain associated with closed TRiC. This indicates that TXD and TXD-CTD are encapsulated upon closure but interact weakly and dynamically with the closed chamber (Figure 5D-ii, iii).

This structural and biochemical analysis reveals the complex role of PhPL2A domains in the interaction with TRiC throughout its ATP-driven conformational cycle. In the open state, it appears TXD and the CTD play key roles in binding and encapsulation. On the other hand, the NTD is an essential component for the proper structural orientation of PhLP2A in the closed TRIC chamber. Our analysis also provides insight into the domain binding and release events leading to PhLP2A rearrangements during the chamber closing process. Each TRiC ring is composed of eight distinct subunits with distinct ATP binding affinities ^7^. As a result, an asymmetric wave of chamber closing has been suggested ^2,7^. Upon closure, each CCT subunit undergoes conformational changes induced by ATP hydrolysis within an allosteric network communicating equatorial and apical domains (Figure 5E-i, ii). We speculate that lid closure releases the NTD from its charged binding sites at the apical domains of CCT3/CCT4, and instead anchors it to the charged surface of the closed chamber at CCT3/6 (Figure 5E-ii, iii). This rearranges and orients PhLP2A within the closed folding chamber. The closure also weakens the open-state interactions of the CTD with CCT3/6, and of the TXD with CCT3 allowing them to be dynamically mobile within the closed chamber. The anchoring effect of the NTD in the closed state may enhance the affinity of TXD and CTD for sites in the chamber, as observed for these domains in the full-length PhLP2A complex with closed TRiC.

### PhLP2A role in TRiC-mediated substrate folding

PhLP2A contributes to TRiC-mediated substrate folding ^23,24^, raising the question of how its distinct domains and interactions affect substrate folding. A recent structure of TRiC directly captured from cell lysate showed that it could form a ternary complex with actin and PhLP2A ^28^. We added PhLP2A to a reconstituted TRiC-mediated actin folding reaction using purified components and a physiologically relevant system ^10^. Briefly, human PFD-bound actin remains in a high protease K sensitive unstructured state (Figure S6A). We used PFD-actin to generate a TRiC-substrate complex followed by the addition of PhLP2A and subsequent incubation with ATP/AlFx (Figure 6A-i). CryoEM analyses revealed most TRiC particles adopted the closed state, with ~42 % of the particles containing extra-density attributable to PhLP2A or actin encapsulated in each different chamber (Figure S6B). Further classification revealed clear actin and PhLP2A density each occupying two opposite chambers (Figure 6B). In the closed TRiC chamber, we identified completely folded actin which indicates overall that unstructured actin was transferred from PFD to TRiC and then subsequently folded by TRiC and PhLP2A upon ATP hydrolysis. Of note, the overall architecture of our structure resembles the previously reported TRiC-actin-PhLP2A structure (Figure 6B) ^28^, which was a single snapshot structure of an undefined folding state captured from a cell lysate. The consistency between these structures indicates our reconstitution strategy successfully mimics physiologically relevant *in vivo* TRiC-mediated folding. Since our result is derived from a folding process reconstituted using purified components, it can provide insight into the folding process resulting from a well-defined chemical reaction. Collectively, our structures allow access into the role of PhLP2A in the complete TRiC-mediated actin folding cycle leading to fully folded actin.

**Figure 6.**
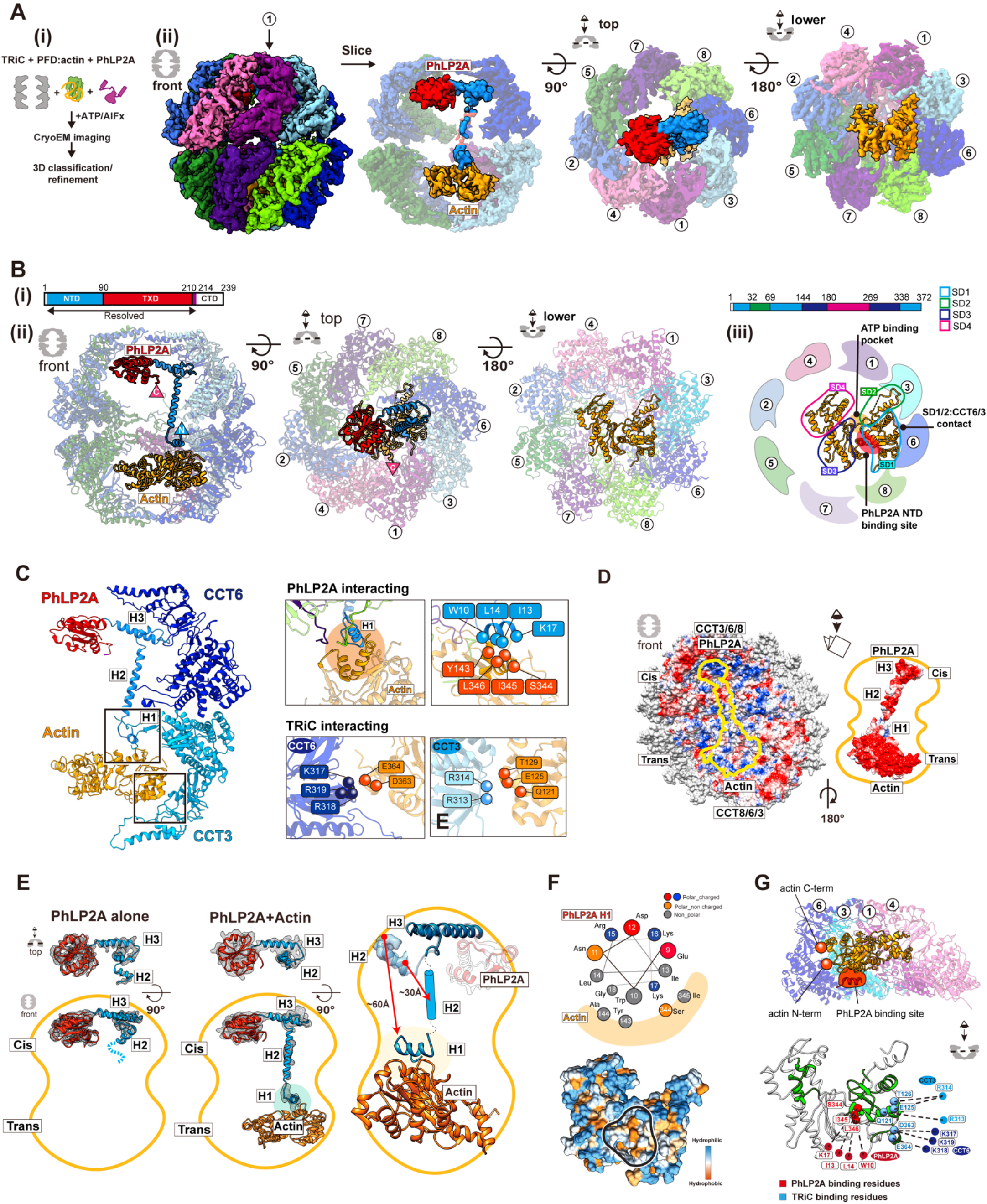
CryoEM structure of closed TRiC with folded actin and PhLP2A occupying each fold ing chamber. (A) CryoEM structure determination of closed TRiC with folded actin and PhLP2A in each chamber. (i) Sample preparation scheme for the substrate-cochaperone encapsulated TRiC. (ii) 3D reconstruction map of folded actin and PhLP2A encapsulated in closed TRiC. The density of H2 of PhLP2A is low-pass filtered and depicted at σ = 1.4 from the density map of the full complex. CCT1 is indicated by an arrow. (B) Atomic model of closed PhLP2A-β-actin-TRiC (i) Schematic of the model of PhLP2A. Modeled domains are color coded. (ii) Front, top, and bottom slice views of PhLP2A and folded actin encapsulated inside the closed TRiC model. (iii) Summary of actin structural features inside the folding chamber. SD1 is the major binding site with the intermediate domain of CCT3/6. The PhLP2A binding site is between SD1 and SD3 (indicated by a red circle), and the ATP binding pocket is on the opposite side, between SD2 and SD4 with fewer constraints (indicated by an orange marker). (C) Detailed interactions between PhLP2A, CCT subunits, and folded actin. (Left) Interactions between the PhLP2A NTD or CCT3 and actin. (Right) (Top) Note that H1 of PhLP2A and CCT3 show direct interaction with actin, forming a local hydrophobic interaction network. Interacting residues are represented as balls. (Bottom) Interacting residues between actin and CCT3/6 are shown as balls. (D) Electrostatic surface of the closed TRiC folding chambers (left), and PhLP2A and actin (right). A symmetric charge distribution inside the closed folding chamber leads to a positively charged surface along the PhLP2A NTD and actin-binding surface. (E) The comparison between PhLP2A with and without actin in the closed folding chamber. Conformational changes of PhLP2A induced by the encapsulated substrate are represented. Once actin is encapsulated in the trans chamber, H1 and H2 of PhLP2A in the cis chamber stretch and traverse the chamber, undergoing a conformational shift of a round ~ 60 Å, leading to direct contact between H1 and actin. TXD and CTD display no change up on substrate encapsulation. (F) The helix wheel plot and hydrophobic surface of encapsulated actin showing the actin-PhLP2A contact sites. (G) (Top) The slice view of the actin-TRiC contact site. CC T4,1,3,6 and actin are shown. N and C terminus of actin are represented as balls and PhLP2A bind ing site is colored in red. (Bottom) The folding defect mutant residues colored in green on the encapsulated actin structure (Rommelaere *et al.,* 2003). Red balls indicate the residues interacting with P hLP2A while blue balls represent CCT interacting residues.

We next examined the molecular contacts between PhLP2A in the cis-chamber and actin in the trans-chamber of closed TRiC. Notably, both PhLP2A and actin interact with the positively charged wall hemisphere comprised of residues in CCT 3/6/8 (Figure 6D). This common binding interface may provide the chemical logic to segregate substrate and cochaperone into two separate chambers. Actin in the chamber appears to be in a fully folded state with its four subdomains (SD1-4) (Figure 6B-ii, iii, Figure S6E). SD1 and SD2 bind through large surface contact with the CCT1/3/6 wall, while SD3 and SD4 extend towards the center of the chamber without direct TRiC interactions. Our structure corresponds to the previous hydrogen-deuterium exchange and mass spectrometry (HDX-MS) experiment of actin in TRiC, which suggested that SD1 and SD2 is the location protected by TRiC ^4^. Notably, the ATP binding pocket is ~20 degrees more open compared to the monomeric actin structure (PDB 4PKH, Figure S6E) ^32^, potentially because of the domain-specific constraints from TRiC. On the other side, the overall architecture of PhLP2A is similar to the one in substrate-free TRiC (Figure 6E, S6F). However, the presence of substrate caused a dramatic change in the NTD H2 to cross the inter-chamber cavity and stretch towards actin (Figure 6E and Movie S4). More importantly, in the presence of substrate, the NTD H1 helix was resolved and shown to be in the trans-chamber making direct contact with folded actin (Figure 6B-ii, 6E, S6F-G and Movie S4). This indicates that the presence of bound actin in the trans-chamber of closed TRiC leads to a ~60 Å substrate-induced conformational change in the NTD of PhLP2A (Figure 6E and Movie S4). The cleft between actin SD1 and SD3 contains a hydrophobic groove which is the site that binds the amphipathic helix H1 of PhLP2A (Figure 6F, S6I and Movie S4). Of note, this hydrophobic groove is an interaction hotspot for regulators of actin polymerization ^33^ as well as a mutation hotspot affecting actin folding (Figure 6G, S6I) ^34^. Taken together, our structural analysis suggests that TRiC and PhLP2A cooperatively orchestrate actin folding. The TRiC chamber wall provides a rigid molecular template to bring the actin subdomains together to establish the topology and H1 of PhLP2A, as an actin specific contact, of PhLP2A serves to seal the exposed hydrophobic surface between SD1 and 3 which may facilitate the formation of this actin lobe and assist folding.

### Conservation of chaperonin and substrate contacts across the PhLP superfamily

Our analysis indicates each domain of PhLP2A plays a singular yet synergistic role in this cochaperone association with and modulation of TRiC. Since PhLPs have diverged into distinct homologs, we next considered the implications of our conclusions to understand the activity of this superfamily and the sources of functional difference ^16,17^. A phylogenetic tree constructed using the full-length sequences of 71 eukaryotic PhLPs revealed the evolutionary hierarchy among PhLP family members (Figure 7A). The simple PhLP system from unicellular organisms (e.g., PLP1 and PLP2 in yeast), diverged into diverse PhLP1-3 subfamilies in multicellular organisms in addition to the phosducin (PDC) family. Human PhLP3 is closest to yeast PLP1 and human PhLP2 is closest to yeast PLP2. We also find the PhLP1-3 branches preceded the appearance of phosducin (PDC), which was historically identified first. PhLP1, 2A and 3 are ubiquitously expressed in a wide variety of tissues ^19,35^ while PDC and PhLP2B are each expressed exclusively in the retina and the reproductive tissues ^16,36,37^. This suggests PDC and PhLP2B diverged later to function in a tissue-specific manner.

**Fig 7.**
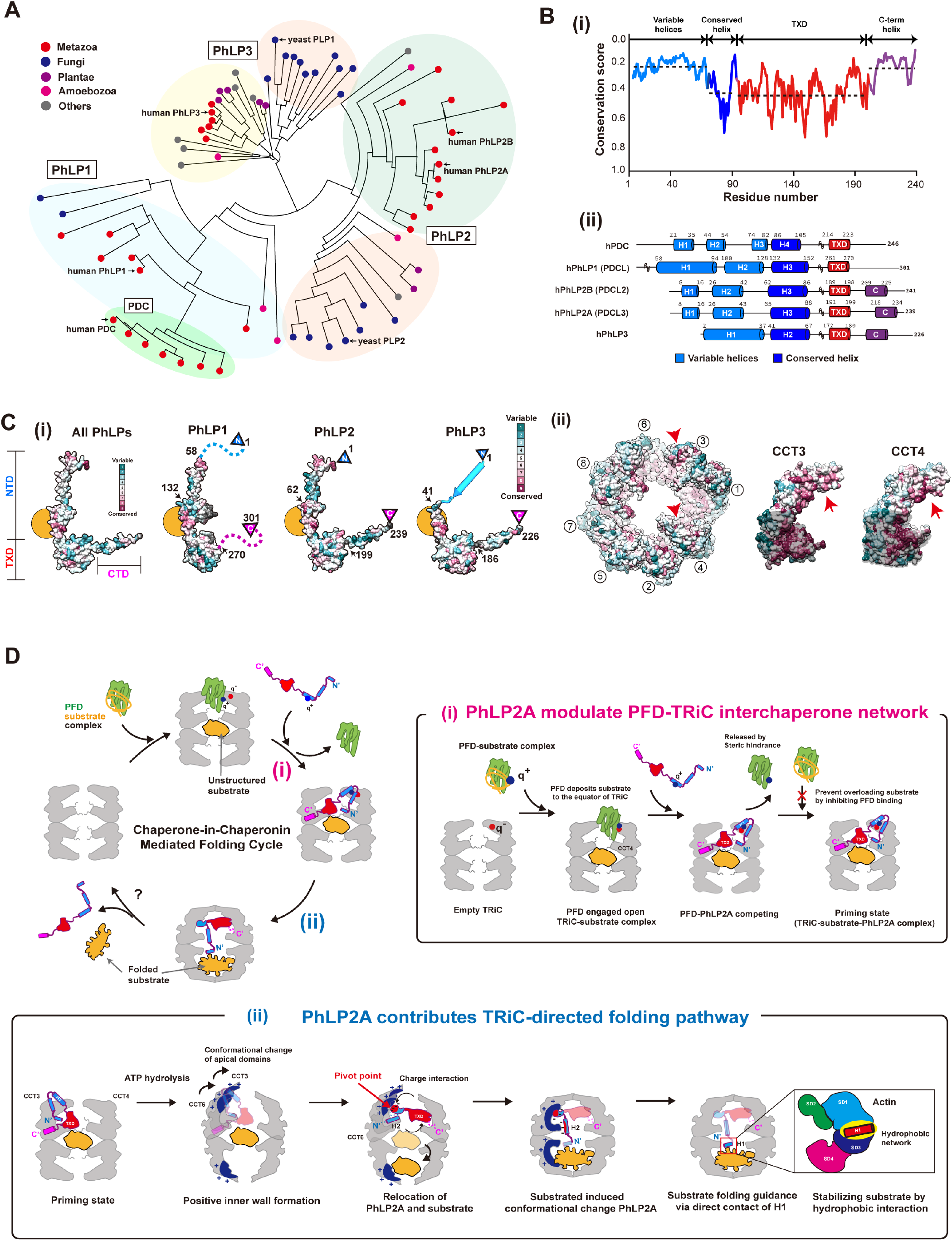
Evolutionary analysis on phosducin-like protein family and the proposed mechanism of chaperone-in-chaperonin mediated folding cycle. (A) Phylogenetic tree of the PhLP family using 71 phosducin-like proteins. Multiple sequence alignment was produced in T-coffee. Colored circles indicate the kingdom of each branch. Each subtype (PhLP1, PhLP2A, PhLP2B, PhLP3, PDC) is grouped and indicated in the phylogenetic tree. (B) (i) The domain-wise conservation score of PhLPs. Each domain is divided according to human PhLP2A dom ain features. The dotted line indicates the average value of the conservation score for each domain. (ii) The secondary structures of 5 human PhLPs based on Alphafold predictions. The cylinder repre sents the predicted helix structure. Residues not shown in the diagram are marked with a tilde. (C) (i) Surface of phosducin family and TRiC colored with conservation scores. (Left) Conservation surfaces of all PhLPs, PhLP1, PhLP2, and PhLP3. All PhLPs shows conservation plot using all 71 phosducin-like protein sequences while the score for each subtype is calculated from the member in the phylogenetic tree. The yellow circle indicates the conserved surface in H3. Arrows indicate the residue numbers which divide the NTD, TXD, and CTD. (ii) Residue conservation among CCT subunits, a nd CCT3, CCT4 is shown. The conservation score is calculated using Consurf and the surface is colored based on conservation scores. Each conservation score is independently calculated and normalized. (D) Proposed mechanism of PhLP2A-mediated substrate folding. At the initial stage of the fol ding cycle, PFD delivers the substrate via direct contact with apo-TRiC. PhLP2A interacts with TRiC and releases PFD. Upon ATP hydrolysis and chamber closing, PhLP2A and the substrate reposition and occupy one chamber each. When PhLP2A makes direct contact with the substrate, substrate fol ding occurs. Finally, TRiC reopens and a folded substrate is released. (i) PhLP2A modulates the PF D-TRiC interaction. PFD captures nascent polypeptides from ribosomes and delivers the substrate by binding directly to empty TRiC. After the full engagement, PFD is stabilized mainly by charge intera ction and deposits the substrate to the inter-ring space of TRiC. When PhLP2A binds to TRiC, its N TD releases PFD by competing with the same binding sites. PhLP2A also inhibits further PFD interactions and possibly prevents substrate overloading. TRiC forms a ternary complex with PhLP2A and the substrate using its three chambers system as a priming state. (ii) PhLP2A contributes to TRiC-mediated substrate folding. When ATP hydrolysis occurs, the positively charged surface inside the TRi C chamber is formed through CCT3/6. The negatively charged NTD of PhLP2A acts as a pivot point to induce 90 degrees relocation of PhLP2A in the cis-ring. The substrate migrates to the trans-ring. Without substrate, PhLP2A is encapsulated in the cis-chamber only. Once the trans chamber is occupied by the substrate, H1 and H2 traverse the two chambers. H2 penetrates the middle of the two rings, and H1 makes direct contact with the substrate. A hydrophobic network between the PhLP2A H1 helix and the substrate might assist the substrate folding. When the chamber opens, the encap sulated components are released.

While all PhLPs share a similar domain architecture, we observe varying levels of conservation in different domains (Figure 7B-i). The domain elements that contact both open and closed TRiC, namely H3 of the NTD and TXD, are highly conserved across PhLPs particularly the TRiC interacting positive patch on H3 and the hydrophobic surface of TXD (Figure 7C, S7A). This pattern of conservation suggests that all PhLP family members interact with open and closed TRiC through similar domain-wise contacts as described here for PhLP2A. Indeed, previous reports indicate that PhLP1-3 can all bind TRiC ^24,36,38,39^. In contrast, the TRiC interacting residues are significantly diverged in PDC (Figure S7A), despite having ~28% sequence identity with PhLP2A; this explains previous findings that PDC does not bind TRiC ^38,40^.

Interestingly, both the NTD and the CTD highly varied among PhLP family members (Figure 7B, S7A-C). For instance, the sequence of PhLP2A contacting TRiC-bound actin in our analysis maps to a variable NTD region (Figure S7B). Such sequence variation may confer PhLPs with the ability to contact different folding substrates within the TRiC chamber. For instance, H1 in the NTD of PhLP2B also has a -WNDIL-actin-binding motif, similar to that identified in PhLP2A in this study and previously ^28^. This sequence is weak in other PhLP variants. On the other hand, the structure of a PDC-Gβ complex identified a -GVI-Gβ binding motif in PDC that is also highly conserved in PhLP1. In contrast, this motif is absent in PhLP2A and PhLP2B, which cannot interact with Gβ ^18^. It thus appears that PhLPs isoforms share conserved TRiC binding elements but evolved distinct substrate-binding elements which determine PhLPs action within the TRiC folding chamber.

## DISCUSSION

### PhLP2A: a cochaperone that functions within the chaperonin chamber

TRiC functions in a cooperative network with cochaperones, including PFD and PhLPs ^14^. While PFD only binds to TRiC in the open state in the absence of ATP ^15^, our analysis shows that PhLP2A can bind to both open and closed TRiC (Figure 1, 4). In both conformations, PhLP2A binds inside the chamber but undergoes significant changes upon TRiC closure and in the presence of substrate (Figure 7D). In the open state, the PhLP2A NTD adopts an extended conformation, with H3 preventing the binding of PFD to TRiC. ATP-dependent TRiC closing induces a large reorientation of PhLP2A domains, with H3 reorienting the NTD and shifting its binding sites within the smaller closed TRiC chamber. Within the closed chamber, PhLP2A becomes compacted with the highly negatively charged H2 and part of H3 associating with the positive hemisphere within the TRiC chamber. The presence of substrate in the opposite closed TRiC chamber induces a further conformational change of PhLP2A, specifically in H1 and H2 of the NTD, whereby H2 traverses the chamber positioning H1 to directly bind the substrate actin. We also find that the PhLP2A CTD is extended from the TXD which proximally locates to the substrate binding site at the inter-chamber space of TRiC. The TXD crosslinks to the flexible CCT tails, which contribute to substrate binding ^10,28,41^. Therefore, it is tempting to speculate that the PhLP2A TXD may sense the substrate directly via the proximal interaction and cascades the binding affinity to the CTD. Overall, PhLP2A functions throughout the different states of TRiC, boosting the detachment of PFD and assisting substrate folding within the chamber. Importantly, the major sites of PhLP2A contact to both open and closed TRiC are conserved across all PhLP family proteins (Figure 7B-C, S7A-B), suggesting our findings may reflect more generally how this family of cochaperones regulates TRiC function.

### Role of PhLP2A in the TRiC-mediated folding cycle

Our results raise intriguing questions regarding the role of PhLPs in the TRIC-mediated folding cycle. Substrates are thought to be delivered to TRiC by Hsp70 and PFD ^42–44^. Chamber closure encapsulates the substrate and promotes folding ^4,10^. There is increasing evidence that the interior of the TRiC chamber plays a multifaceted and complex role directly in the folding of the bound substrate. In the open state, the combinatorial interaction with diverse apical domain binding sites influences the topology of the bound substrate ^4,5^. In addition, the flexible CCT N- and C-termini at the equator of the chamber also interact extensively with the substrate. Upon lid closure, the encapsulated substrate interacts extensively with the closed wall as well as the N- and C-tails. In this context, the fact that PhLP2A domains bind to multiple sites inside the chamber in both open and closed TRiC states is remarkable, particularly in light of recent findings showing interactions with the TRiC chamber direct the folding trajectories of at least some substrates such as tubulin and actin ^4,10^. Our findings suggest that PhLPs must modulate the interactions and/or conformation of the TRiC-bound substrate inside the chamber, and possibly the TRiC conformational cycle itself. Future biophysical and structural experiments will be required to ascertain how PhLPs modulate TRiC-mediated folding of the substrate. Nonetheless, based on our structural analyses and previous experiments we can speculate on possible aspects of the interplay between TRiC and the substrate. Our finding that PhLP2A H3 shares a TRiC binding site with PFD and their binding is mutually exclusive (Figure 3, S3A, S3F) suggests PhLP2A coordinates association between PFD and TRiC. Previous *in vitro* translation reactions in rabbit reticulocyte lysate showed the addition of high levels of PhLP3 led to the accumulation of actin-bound PFD ^24^ and inhibited TRiC-mediated folding of newly synthesized β-actin ^38^. Perhaps forcing high amounts of PhLPs into this system saturates the apical domain substrate-binding sites in the TRiC chamber precluding substrate binding. How these observations relate to the function of PhLPs in the TRiC-mediated folding reaction remains to be investigated.

Our analysis of the interaction of PhLP2A with actin enclosed within the closed chamber of TRiC provides another clue as to the possible role of PhLP2A in modulating folding intermediates within the closed TRiC chamber. Thus, we observe the H1 of PhLP2A forms a buried hydrophobic core with helix 340-350 of encapsulated actin (Figure 6F, Figure S6H). Our data suggest PhLP2A stabilizes an exposed hydrophobic core of actin intermediates that are folding within the TRiC chamber. While future experiments should delve into the mechanism of how PhLP2A assists folding, it is noteworthy that mutation in actin residues 340-344, *i.e.* corresponding to the PhLP2A H1 binding site, significantly affects actin folding ^34,45^. Furthermore, previous studies observed that the folding of SD1 is rate-limiting for TRiC-mediated actin folding (Figure 6G) ^4^. Taken together, our study provides mechanistic insight into the molecular interplay between TRiC and PhLP2A to orchestrate a hydrophobic collapse that drives the folding of the substrate.

### TRiC-PhLPs expand repertoires with substrate-specific regulation

TRiC facilitates the folding of ~10% of the proteome and diversifies the strategies to support folding for many different types of protein substrates. The heterooligomeric nature of TRiC is considered key to diversifying the recognition of distinct motifs in its diverse substrates ^4,5,46^, as well as promoting their folding within the closed chamber ^4,10^. Our analysis of PhLP family evolution suggests the modality of binding of family members to the TRiC chamber is largely conserved, whereas the substrate binding elements within the NTD, and perhaps the CTD, have diversified to assist folding to different substrate proteins. Indeed, different PhLP family members have different substrate specificity and function ^21,22^ through substrate-specific sequences in highly diverged NTD ^23,24^. While this hypothesis will require direct experimental testing, we propose PhLPs assist TRiC to support the folding of different subsets of proteins.

PhLPs are only found in eukaryotes and their advent may be linked to the increase in proteomic complexity ^47^. We propose that the combinatorial usage of PhLPs as substrate-specific cochaperones expands the ability of TRiC to fold different substrates. Notably, the activity of PhLP can be regulated by phosphorylation ^48 49^, adding another layer of regulation to TRiC action in the cell. Our work uncovers another layer of complexity in the regulation of TRiC cochaperone networks mediating cellular folding.

## ACKNOWLEDGMENTS

This work has been supported by the National Institutes of Health grant R01GM074074 to JF and the Korean National Research Foundation (2019R1C1C1004598, 2020R1A5A1018081, 2021M3A9I4021220, 2019M3E5D6063871) and SUHF foundation to S.-H.R. The cryoEM data was collected and processed at The Center for Macromolecular and Cell Imaging (CMCI), Institute for Basic Science (IBS), Global Science experimental Data hub Center (GSDC) at Korea Institute of Science facilities. We thank Dr. Bum Han Ryu (Institute for Basic Science (IBS), Korea) for supporting cryoEM data collection. We also thank Dr. M. Steinegger and Dongwook Kim from Steinegger lab for discussions on evolutionary analysis. We thank members of CMCI for their discussions and suggestions. We thank members of the Frydman lab for useful discussions and suggestions. We thank Dr. P. Picotti and the UCSF Mass Spectrometry Facility for access to instrumentation.

## AUTHOR CONTRIBUTIONS

Conceptualization, J.F., D.G. and S.-H.R.; Investigation, D.G., J.P, H.K. A.L. and S.L.; Writing, J.F., D.G., J.P., H.K., S.L. and S.-H.R.; Funding Acquisition, J.F. and S.-H.R.

